# Genetic encoding of 3-cyano-tyrosine and its use in controlling the chromophore isomeric state of the fluorescent protein mKate

**DOI:** 10.64898/2026.07.16.738936

**Authors:** Connor J. Stevenson, John J.K. McLarnon, James Harnedy, Salma A. Elsherbeni, Debarshi Saha, Wolfgang Langbein, Paola Borri, Jamie A. Platts, Louis C Morril, D. Dafydd Jones

**Affiliations:** Molecular Bioscience, School of Biosciences, Cardiff University, Sir Martin Evans Building, Cardiff CF10 3AX, UK; School of Chemistry, Cardiff University, Cardiff CF10 3AT, UK; Department of Chemistry, University of Bath, Bath BA2 7AY, UK; School of Physics and Astronomy, Cardiff University, Cardiff CF24 3AA, UK

**Author notes:** These authors contributed equally (Connor J. Stevenson, John J.K. McLarnon).

## Abstract

Switchable β-barrel-type fluorescent proteins are essential genetically encoded probes for super-resolution imaging. The space required for chromophore *cis–trans* isomerisation can also provide an opportunity to introduce bulkier chemistry at the 3-position of the phenolic ring. Here, we report, to our knowledge, the first successful genetic encoding of 3-cyano-L-tyrosine (3CNY) into a protein. Using genetic code expansion, the cyano-containing amino acid was incorporated directly into the chromophore of mKate, a pH-dependent switchable red fluorescent protein. In mKate, the chromophore adopts a fluorescent phenolate *cis* state at physiological pH, transitioning to a phenolic *trans* state under acidic conditions. Substitution of the native tyrosine with 3CNY yields a functional protein exhibiting hypsochromically shifted spectral properties. Time-dependent density functional theory (TD-DFT) calculations indicate that 3CNY incorporation results in a *trans* state at pH 8. Unlike mKate, the *trans* state is fluorescent. In contrast, incorporation of 3-chloro-L-tyrosine (3ClY) preserves the preference for the *cis* phenolate state. Molecular modelling suggests that the cyano group can form stabilising hydrogen bonds with residues S143 and S158, promoting the *trans* configuration. DFT analysis further indicates that the electron-withdrawing cyano group perturbs conjugation across the chromophore, potentially lowering the barrier to *cis–trans* isomerisation. Conversely, wild-type and 3ClY variants maintain polarised HOMO and LUMO distributions in the *cis* state, supporting stronger conjugation and a reduced HOMO–LUMO gap. Overall, the introduction of a genetically encoded 3-CNY tyrosine analogue into a fluorescent protein chromophore expands our mechanistic understanding and enables incorporation of a new chemical tag directly into the chromophore.

## 1. Introduction

Classical β-barrel fluorescent proteins (FPs) represent an important class of proteins due to their role as genetically encodable imaging tags [1–3]. The modern repertoire of FPs spans the blue to far red region acting as both passive and active imaging tags,[2,4] and even ventures into the realms of nanoscience [5–7]. The chromophore is central to FP function and is encoded directly within the amino acid sequence. Three contiguous residues, Xaa-Tyr-Gly (where Xaa is variable), enclosed by the β-barrel structure comprise the chromophore. In the presence of O_2_, the three residues undergo covalent rearrangement to form the chromophore (Figure 1a), with the phenol group (P ring) of the tyrosine linked to the newly formed imidazolinone (I ring) via a β-methylene bridge [8,9]. For red FPs, an additional oxidation event occurs forming an N-acylimine so extending the conjugated network into the residue preceding Xaa [10].

**Figure 1.**
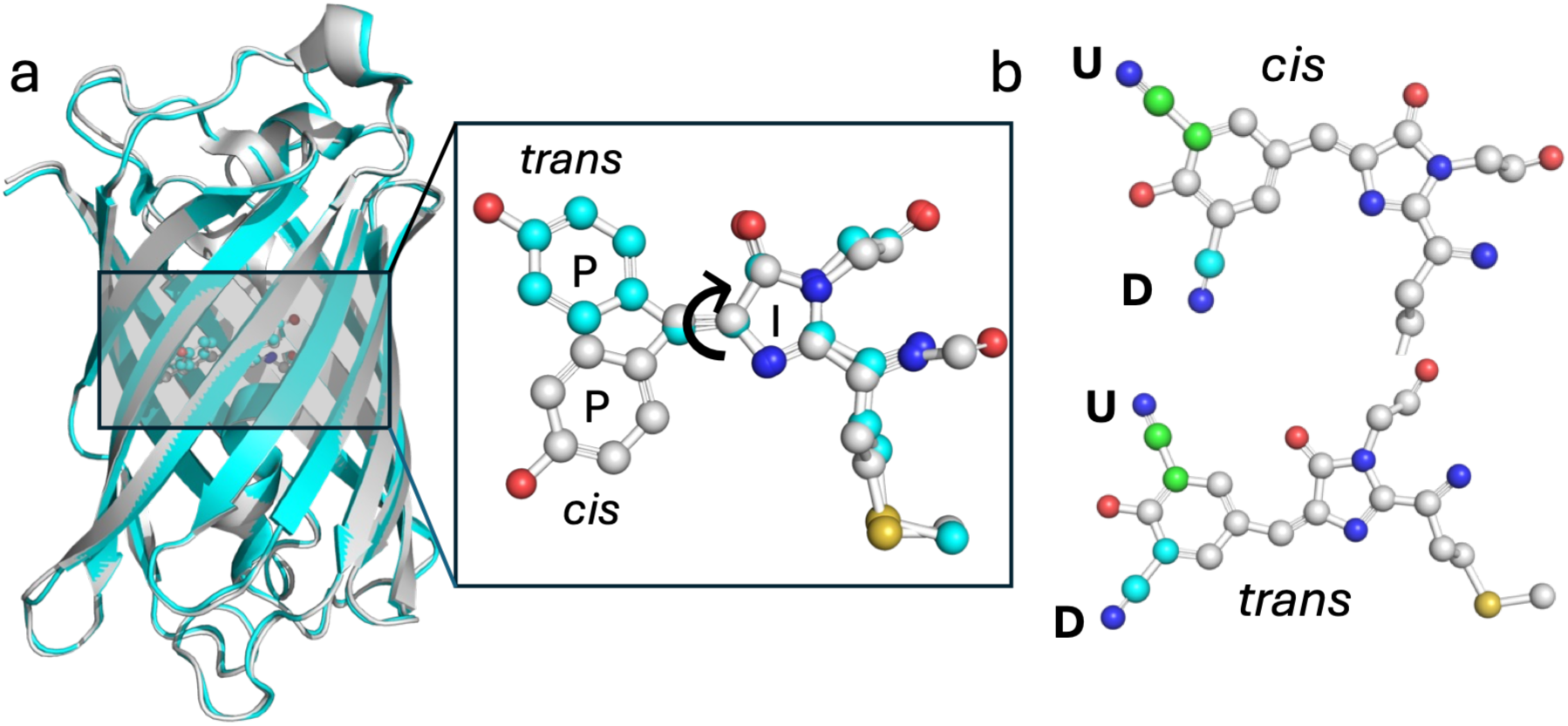
Chromophore structure of mKate. (a) The β-barrel and chromophore structures of mKate at pH 7 (grey, PDB 3BXB) and at pH 4.2 (cyan, PDB 3bx9) [11]. Inset are the two alternative chromophore states with the P and I rings are labelled for the *cis* (grey carbons) and *trans* (cyan carbons) states. (b) Putative models of the 3CNY-derived mKate chromophores. The **U** and **D** notation refers to which rotamer state the cyano group occupies; the **U** state carbons are coloured green, and the **D** state carbons coloured cyan. Note, that only either the **U** or **D** form is sampled at any one time but are both shown within the same chromophore structure for simplicity.

The P and I rings can potentially exist as *cis* or *trans* isomers centred around the β-methylene C=C (Figure 1a). The majority of FPs form the cis isomer as their fluorescence permissive state. A few rare cases such as eqFP611 [12] and its engineered derivatives such as mRuby [13] exist in the *trans* isomeric fluorescent state. Others can switch between the two states, with each state having discrete spectral properties. Such switchable FPs have proved important probes for high-resolution imaging [14] whereby fluorescent proteins can be “switched on” post-translationally and post-maturation when and where required [15,16]. Switching is most commonly induced photochemically, as is the case for FPs such as Dronpa [17], Kaede [18] and the EosFP family [19]. Photoswitching occurs through the application of certain wavelengths of light. For example, Dronpa switches from its dark state to its fluorescent state on application of UV-violet light (∼400 nm) and converts to the dark state on irradiation with light (488 nm) close to its λ_max_ (503 nm) [17]. Generally, the *cis* state is the most red shifted and brightest form. Other FPs, such as pHlourin [20] and mKate [21] exist predominantly in the *cis* forms at physiological pH but convert to the *trans* state under acidic conditions [11]. The mechanism is thought to involve protonation of the phenol hydroxyl group, with the *cis* form favouring the phenolate and *trans* the phenolic form. As with the photochemically switchable FPs, the *trans* state is generally blue shifted compared to the *cis* state.

For the chromophore to sample both isomeric states, space within the chromophore pocket must be available to accommodate both forms and allow rotation of the phenol ring. For example, both isomers in the crystal structure of mKate at pH 4.2 are observed [11]. Ergo, space is available to incorporate new chemical entities at the 3 (*meta*) position of the P ring through incorporation of tyrosine derivatives using an expanded genetic code [22–24]. Genetic code expansion enables new chemistry to be incorporated into a protein, so expanding its functional landscape [25]. Many genetically encodable non-canonical amino acids (ncAAs) are based on tyrosine [25] and thus can theoretically be incorporated directly into the FP chromophores. Incorporation of ncAAs with the tyrosyl OH replaced by another chemical entity is relatively common and has been applied to FPs, resulting in new versions with useful spectral properties such as photo-activation [26–30]. However, the phenolic/phenolate (Ph-OH/Ph-O^-^) group and transition is critical to the spectral properties of FPs [1,8]; replacing the OH with another group (e.g. azide, nitro, cyano) results in a blue shifted chromophore with poorer spectral properties. Alternatively, the new chemical group can be placed at the 3-position but adds steric bulk within the tight confines of the buried chromophore pocket [22,23,31]. While halo-tyrosine ncAAs have been successfully incorporated into FPs [24,32] larger groups such as NO_2_ (3-nitro-tyrosine) are more disruptive, especially in terms of the impact on fluorescence properties. [24,33]. Boxer and colleagues addressed this problem by using the photo-switchable FP Dronpa2 and a circular permutated version of GFP. They successfully incorporated a range of ncAA version of tyrosine including 3-OCH_3_, 3-CH_3_, as well as 3-NO_2_ [24].

The cyano (-C≡N) group, also commonly known as a nitrile group, represents a potentially useful chemical entity to incorporate into the chromophore of a protein, especially for alternative, vibrational-based imaging approaches such a stimulated Raman spectroscopy (SRS) [34–36]. Cyano groups generate spectrally narrow Raman resonances in the biologically silent vibrational window. However, spontaneous Raman scattering produces weak signals as compared to fluorescence. By coupling a cyano group to electronically active synthetic dyes, electronic pre-resonance SRS increased signal strength and in turn sensitivity by orders of magnitudes, making cellular imaging a possibility [37]. To rival fluorescence microscopy, such SRS-based approaches need to enable genetic encoding of suitable probes [38]. Incorporating a cyano group in place of the chromophore’s phenolic OH group has been possible through incorporation of 4-cyano-L-phenylalanine but results in hypsochromically shifted spectrally compromised FPs [26].

Here, we describe the synthesis of 3-cyano-L-tyrosine (3CNY) and demonstrate its ability to be genetically encoded into proteins, including within the chromophore of the pH switching RFP mKate. We show that the incorporation of 3CNY is tolerated within mKate’s chromophore and results in a blue shifted conformationally restricted form of the mKate chromophore that we suggest occupies dominantly the *trans* isomeric state. Density functional theory indicates that the electron withdrawing nature of the cyano group reduces conjugation across the chromophore system, potentially lowering the barrier for *cis-trans* isomerisation; new H-bonds between the cyano group and the protein then potentially stabilise the *trans* state.

## 2. Results and discussion

### 2.1 Synthesis and incorporation of 3CNY

As 3CNY (**4** in Figure 2) is not commercially available, we synthesised the ncAA over 3 steps from commercially available 3-iodo-L-tyrosine (**1** in Figure 2). Amino acid **1** was protected by reaction with 9-borabicyclononane (9-BBN) to form boroxazolidinone **2** with a 81% yield,[39] which was used in a Pd-catalysed cyanation to access protected 3-cyano-L-tyrosine **3** in 42% isolated yield after chromatographic purification *via* silica gel flash chromatography.[40] Deprotection by treatment with MeOH:CHCl_3_ (1:7.5) at room temperature for 16 h afforded the desired 3CNY **4** in 78% yield after concentration *in vacuo* without the need for further purification.[41] Synthesis of 3CNY **4** was confirmed by ^1^H NMR (Supplementary Figure S1), ^13^C NMR (Supplementary Figure S2) and mass spectrometry (Supplementary Figure S3).

**Figure 2.**
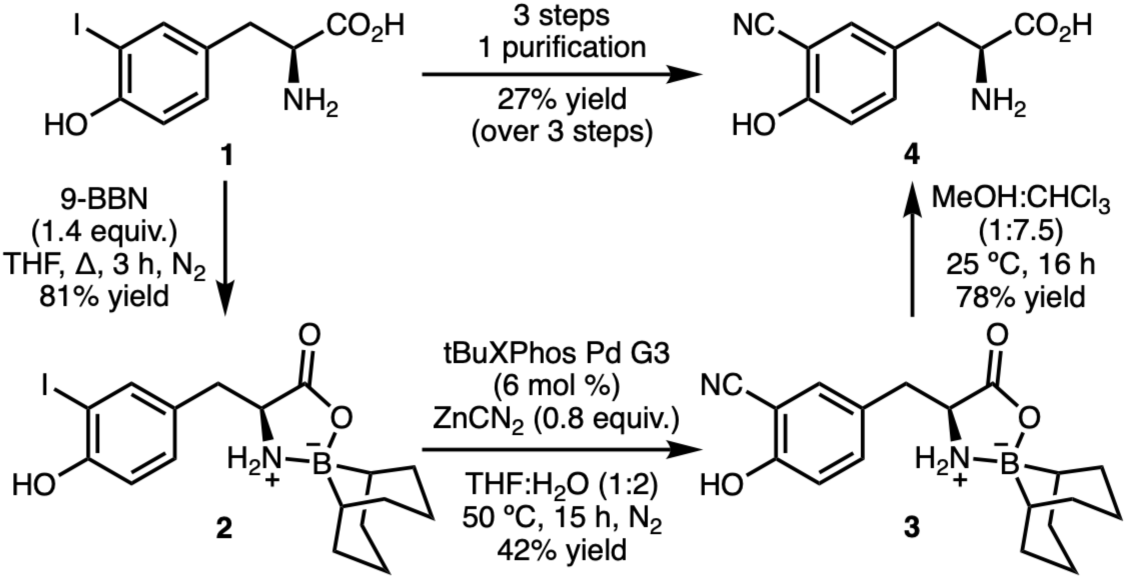
Scheme for the synthesis of 3CNY **4** from 3-iodo-L-tyrosine **1.**

To incorporate 3CNY we first tested an existing amino-acyl tRNA synthase(aaS)/tRNA pair engineered to incorporate 3-nitro-L-tyrosine [22]. Given the general promiscuity of engineered aaS/tRNA pairs [42,43], we surmised that the binding pocket for the nitro group could also accommodate the nitrile group. We used the superfolder GFP [44] variant with the codon for Q204 replaced with TAG as this position is known to incorporate a wide range of different ncAAs [6,7,45,46]. Functional full-length protein was only produced in the presence of 3CNY and mass spectroscopy confirmed incorporation (Supplementary Figure S4).

### 2.2 Comparison of experimental versus DFT predicted spectral characteristics of mKate

The chromophore of mKate switches between a *cis* (or Z) isomeric state at physiological pH to the *trans* (or E) state in acidic conditions. Density functional theory (DFT) can be used to predict the optical properties of FPs [47–51] and inform us of the most likely structure formed by each variant. To test this, we compared experimentally observed electronic excitation data (absorbance spectra) with DFT analysis for the 4 different forms of the mKate chromophore: *cis* or *trans* state that sample either the phenolic (neutral Ph-OH) or the phenolate (anionic Ph-O^-^) form (see Supplementary Table S1).

At pH 8, the observed λ_max_ is at 588 nm with a shoulder around 550 nm (Figure 3a). The closest wavelength from time-dependent DFT (TD-DFT) corresponds to *cis* phenolate form (see Supplementary Table S1), which has predicted wavelengths of 565 nm (major) and 526 nm (minor) (Figure 4b). This tallies with previous observations, including crystal structures, that the *cis* state is present at higher pH values [11]. The deprotonated phenolate form is generally red shifted compared the protonated forms as the phenolate is a stronger electron donor that increases the number of resonance structures and so promotes electron delocalisation across the chromophore. The *cis*-phenolate form is also fluorescent, emitting with at a maximal wavelength (λ_EM_) of 622 nm, a quantum yield (QY) of 33% and brightness of 15.2 mM^-1^cm^-1^ (molar absorbance 46 mM^-1^cm^-1^) [11,21].

**Figure 3.**
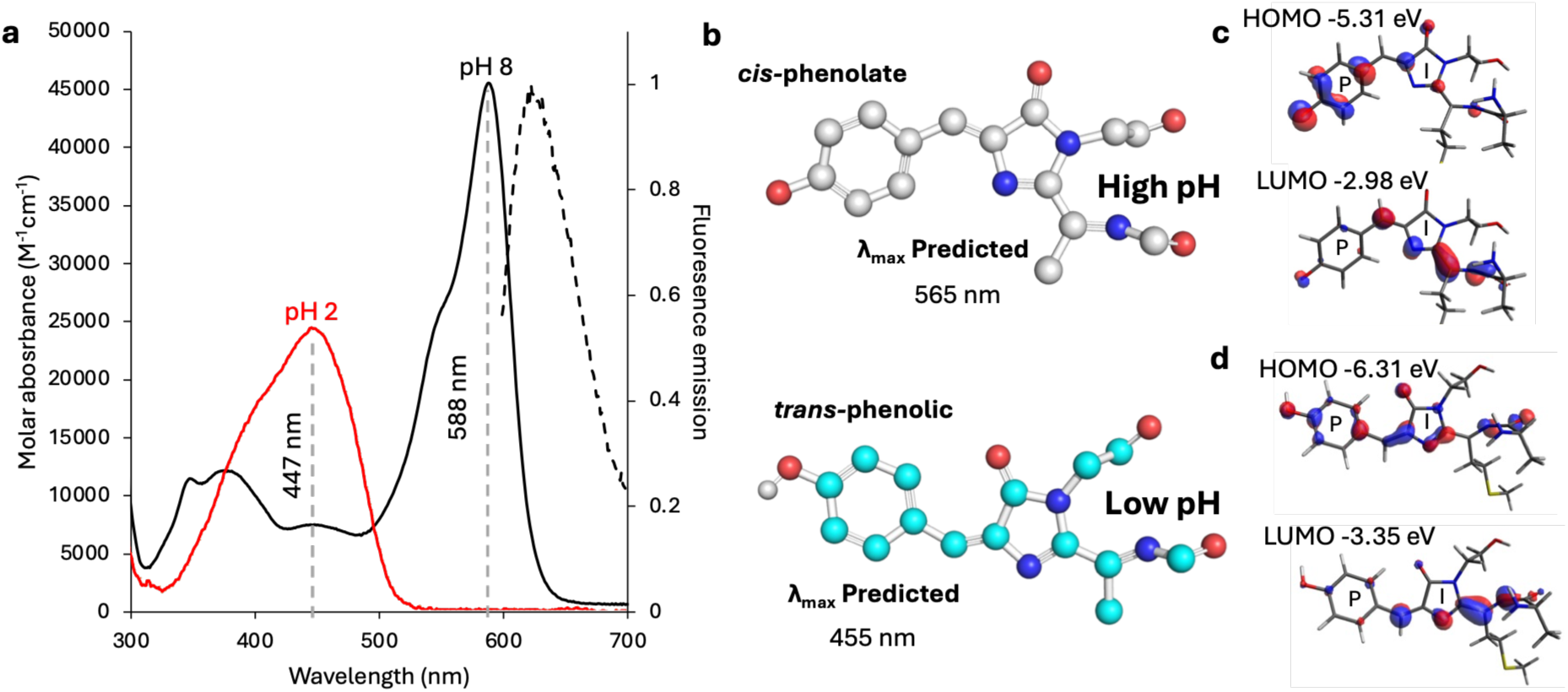
Effect of pH on the spectral and chromophore properties of mKate. (a) Absorbance (solid line) spectra of mKate at pH 2 (red) and pH 8 (black). Fluorescence emission spectra on excitation at 589 nm (black dashed line). No fluorescence emission was detected on excitation at 447 nm. (b) The predicted chemical forms for the mKate chromophore at high pH (grey carbons) and low pH (cyan carbons). The predicted λ_max_ values are taken from Supplementary Table S1. Frontier molecular orbitals of (c) the *cis* phenolate and (d) *trans* phenolic mKate chromophore. The highest occupied molecular orbital (HOMO) and lowest unoccupied molecular orbital (LUMO) forms are shown as isosurfaces.

**Figure 4.**
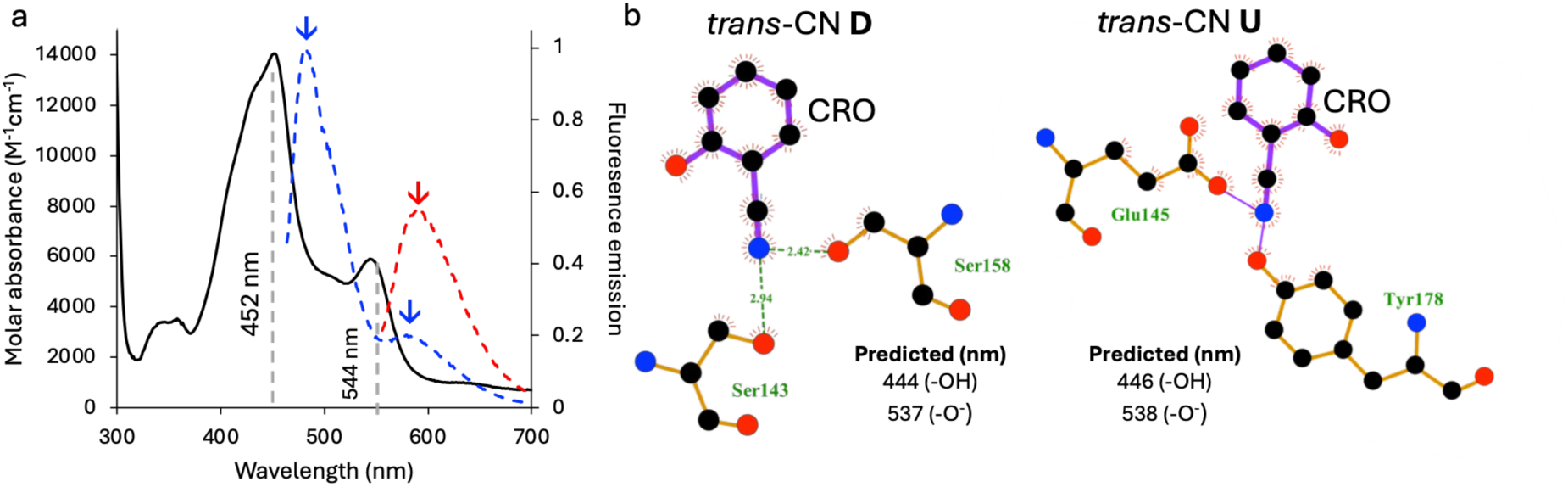
Incorporation of 3CNY into mKate’s chromophore. (a) absorbance (solid black line) and fluorescence emission on excitation at 454 nm (dashed blue line) and 546 nm (dashed red line). The two λ_max_ are annotated. The fluorescence emission is normalised to the peak maximum on excitation at 454 nm. The downwards arrows indicate fluorescence emission peaks. (b) Ligplot+ analysis of the chromophore environment with the predicted values from DFT shown. Dashed green lines between residues (orange bonds) and the chromophore (purple bonds) indicate H-bonds, with the distances shown in Å; solid thin purple lines indicate potential covalent bonds, according to LigPlot+. The predicted peak absorbance values based on DFT are also shown (see Supplementary Table S1).

From DFT analysis, deprotonated Ph-O^-^ forms of the mKate chromophore has differences between *cis* and *trans*: the *cis* form shows significantly red-shifted absorbance compared to the *trans*. In both cases the strongest absorbance band is dominated by HOMO to LUMO frontier orbitals (Figure 3b). Deprotonation of phenolic group raises the energy of both HOMO and LUMO, but the localisation of the former is on the P ring and the latter on the I ring/N-acylimine system. This is raised markedly more for the P ring (∼1 eV) than the I ring/N-acylimine (∼0.3 eV), and hence notably lower energy gap for electronic excitation. The shift in electron density from the P to I ring suggests an intramolecular charge transfer (ICT) process. The phenolate P ring acts an electron donor while the I ring/N-acylimine is the electron acceptor forming a “push-pull” conjugated system. The implications are an enhanced conjugation system resulting in a smaller HOMO-LUMO gap and thus a red shift in fluorescence. ICT is commonly associated with sensitivity to processes such as pH, [52,53] as is the case for mKate.

At pH2, mKate still absorbs visible light but has no detectable fluorescent. The main absorbance peak is broad, blue shifted with a maximum at 447 nm, and has a reduced molar absorbance compared to pH 8 (Figure 3a). From TD-DFT, the chromophore form with the closest λ_max_ is *trans* phenolic (*trans*-Ph-OH in supplementary Table S1 and Figure 3b), with a value of 455 nm, slightly closer to experimental values than the *cis* phenolic form (*cis*-Ph-OH) at 462 nm. Given the evidence available from the crystal structure (Figure 1b [11]) at low pH, the *trans*-phenolic form is the most likely form sampled. DFT indicates that *cis-trans* isomerism of the mKate chromophore does not strongly affect the electronic structure or predicted absorption spectrum of the protonated OH form (Supplementary Table S1). Frontier orbitals are very similar in energy, as is the energy gap between them. HOMO frontier orbitals are spread across the conjugated bond system, while LUMO is located more on the I ring and linked N-acylimine. Thus, in the *trans*-phenolic system, ITC plays less of a role with *π*-*π*\* transitions becoming more significant.

Given the known limitations of standard functionals and finite basis sets in TD-DFT calculations, as well as the fact that these only used the chromophore, omitting the rest of the protein and water that are known to influence the fine spectral properties of FPs [47,49,54], the agreement with experiment noted above is likely to arise from some error cancellation. Nevertheless, DFT is thus a good way of assigning different isomeric forms sampled based on comparison with observed spectral properties.

### 2.3 Incorporation ncAAs into the chromophore of mKate

We next assessed the ability of mKate to tolerate the incorporation of 3CNY within the chromophore. Incorporation of 3CNY within the chromophore can lead to the cyano group potentially occupying two atropisomeric rotamer forms, termed here the **U** and **D** forms (Figure 1b), as well as the *cis*-*trans* and phenolic/phenolate forms. Simple modelling was undertaken using the low and high pH crystal structures to ascertain which rotamer forms can be introduced into the chromophore without generating steric clashes. On analysis of the models using MolProbity [55], the *cis*-**D** had some potentially significant clashes with L199 (Supplementary Figure S5).

Incorporation of 3CNY in place of the mKate chromophore forming residue Y64 (via amber stop codon reprogramming [31]) generated a functional, fluorescent protein (termed here mKate-CRO-CNY) that is spectrally different compared to mKate (Figure 4a). Two peaks are observed at pH 8, a major peak at 452 nm and a minor peak at 544 nm. Based on the TD-DFT analysis, the 544 nm peak is likely due to *trans*-phenolate form (*trans* 3-CN **X** Ph-O^-^) that has a predicted λ_max_ of 537 and 538 nm for the **D** and **U** forms, respectively. In the *trans* phenolate form, both HOMO and LUMO are lowered in energy compared to mKate but here the HOMO is reduced slightly more, giving a wider energy gap (Figure 5a), resulting in the blue shift of the main absorbance band. As with the native mKate chromophore, the HOMO electron density is largely centred on the P ring, whereas the LUMO has more I ring/ N-acylimine character (Figure 5a), suggesting absorbance has similar ICT nature as mKate. The presence of the electron withdrawing 3-cyano group however reduces density of the P ring in the HOMO (compare Figure 5a with Figure 3c) so reducing donor strength of the phenolate compared to mKate and less able to support the delocalised negative charge, resulting in the increased HOMO-LUMO energy gap. The lower electron density on the HOMO P ring may result in a more directional P ring to I ring charge transfer and could make the chromophore more sensitive solvent effects and protonation.

**Figure 5.**
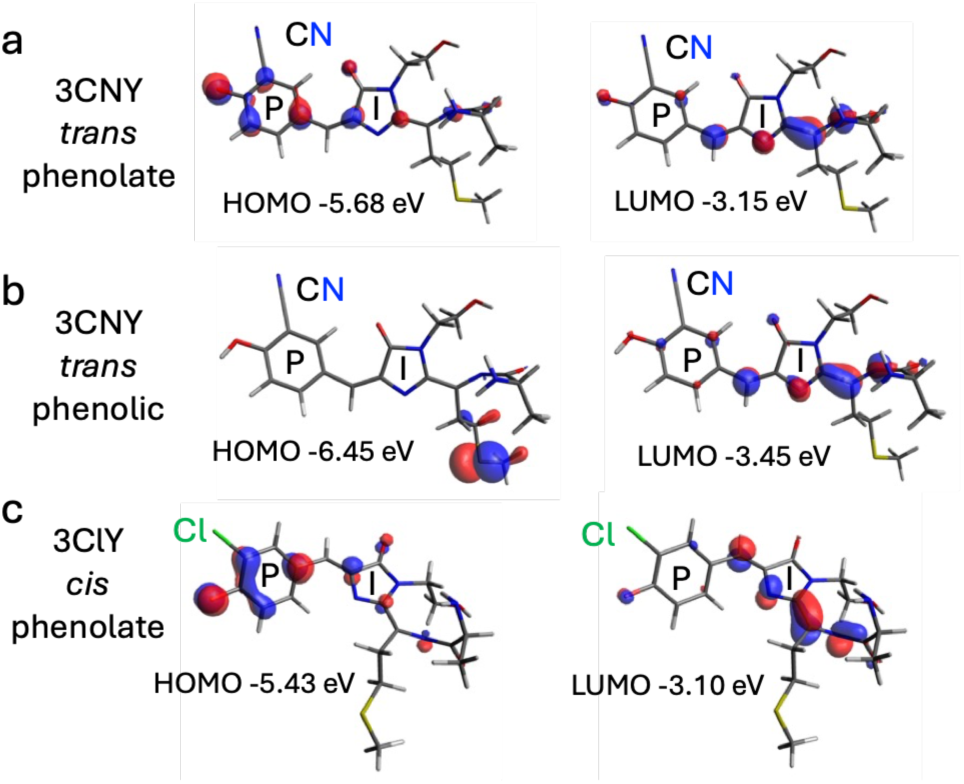
Frontier molecular orbitals of ncAA versions of the mKate chromophore. The HOMO and LUMO forms of mKate-CRO-3CNY in the (a) *trans* phenolate and (b) *trans* phenolic together with (c) the mKate-CRO-3ClY *cis* phenolate

**Figure 5.**
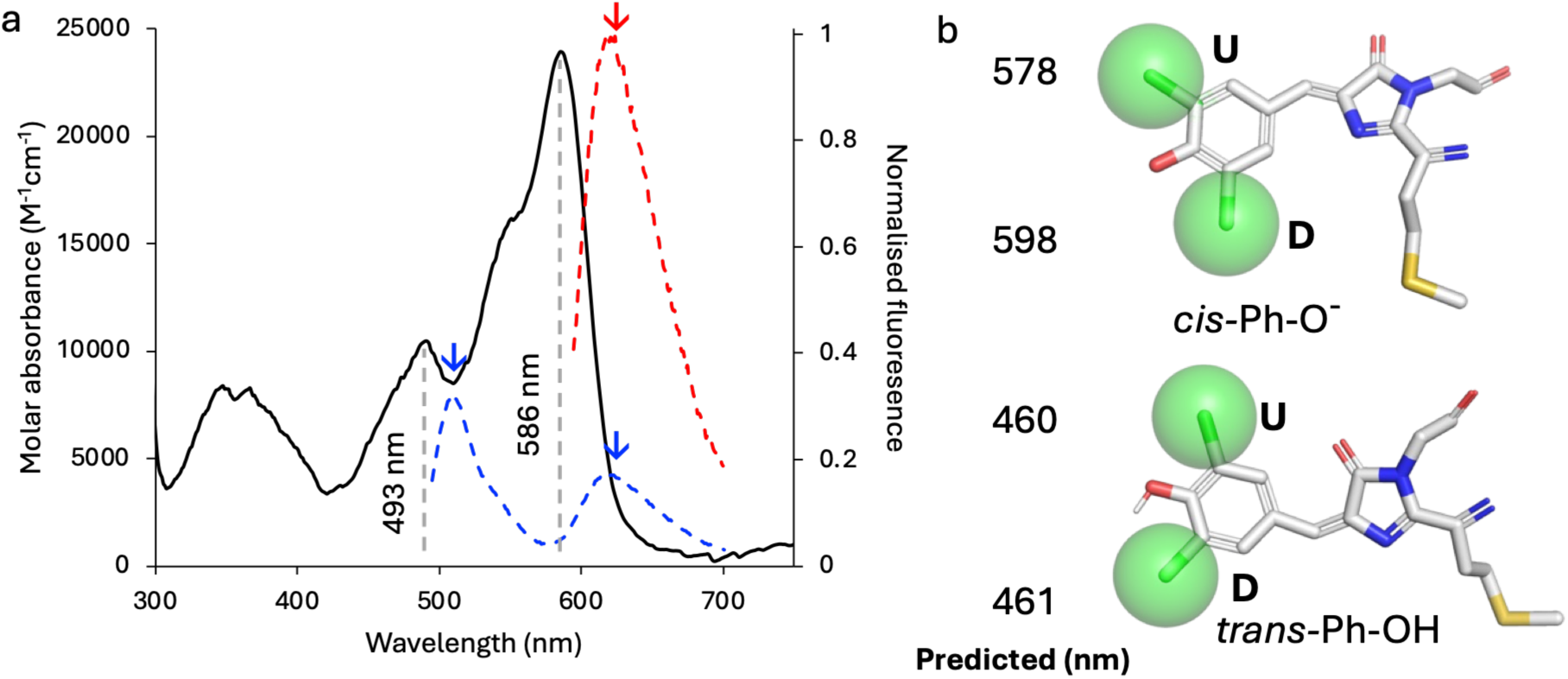
Incorporation of 3ClY into the mKate chromophore. (a) Absorbance spectrum of mKate-CRO-ClY (black line) and emission spectra on excitation at 485 nm (blue dashed line) or 585 nm (dashed red line). Fluorescence emission is normalised to the peak value on excitation at 585 nm. The downwards arrows indicate fluorescence emission peaks. (b) The potential isomeric forms of 3ClY in the mKate chromophore together with their predicted DFT peak absorbance. The green spheres represent the sites of the chlorine atom if either the **U** or **D** forms are sampled.

Excitation of the 454 nm peak species results in a dual emission profile with a major emission peak at 480 nm and less intense peak at 579 nm. While the origin of the secondary fluorescence emission peak is unknown, we speculate that excited state deprotonation or, more commonly known, as excited state proton transfer (ESPT) could partially be occurring here [8,56,57]. ESPT is observed in the original GFP from *Aequorea victoria*, in which the ground state is dominated by the phenolic chromophore but emission is primarily from the phenolate chromophore [1]. This results in a large Stokes shift (∼100 nm) between the ground state absorbance and excited state emission wavelength. However, it is suggested here that the mKate-CRO-CNY emissive form is chemically similar in part to the ground state (i.e. the phenolic form) due to the major emission peak having a relatively small Stokes shift (33 nm). The 452 nm peak likely results from the protonated, *trans-*phenolic form (*trans* 3-CN **X** Ph-OH), which is based on the TD-DFT predicted wavelengths of 444 nm and 446 nm for the **D** and **U** forms, respectively (Supplementary Table S1). Unlike mKate, mKate-CRO-CNY does not show any pH responsive switching (Supplementary Figure S6). At pH 4, the absorbance spectra is too noisy to discern any clear peaks; fluorescence emission on excitation at either peak absorbance wavelength drops. This suggests mKate-CRO-CNY is structurally more sensitive to pH than the original mKate; rather than switching chromophore states, lowering pH could lead to significant structural perturbation or even unfolding.

While DFT cannot not differentiate between the **D** and **U** forms, we propose that the *trans* 3-CN **D** form is most likely to form. Analysis by LigPlot+ [58] suggests that for both rotameric forms, the cyano group will make contacts with the rest of the protein (Figure4b). The **U** rotamer will potentially make what LigPlot+ classes as covalent bonds with the carboxyl group E145 and the hydroxyl group of Y178. As it unlikely such covalent bonds are formed, it suggests that these groups are very close, if not sterically prohibitive, so possibly hindering this conformation. In the **D** form, the cyano group is modelled to make H-bonds with S143 and S158. In mKate, S143 is thought to play a role in stabilising the *cis* state by forming an H-bond with the phenolate oxygen [11]; if S143 is trapped in a H-bond with the 3-cyano group, this may promote the *trans* form and hinder ground state deprotonation. Previous structural studies on GFP-type proteins suggests that the *cis* isomer is formed with the equivalent of the **U** rotamer observed for the majority of the 3-tyrosine derived variants, including nitro and O-methyl [24,33]; the caveat is that in the *cis* isomer, the cyano group would occupy a similar spatial position in the **U** form as it would in the **D** form for the *trans* isomer (Supplementary Figure S7).

In the phenolic form, the HOMO exhibits minimal electron density across the conjugated backbone, with a drop in energy (Figure 5b), reflecting loss of phenolate-driven π-donation, while the LUMO remains localized on the I ring/N-acylimine, indicating a transition to an electronically depleted, acceptor-dominated excited state with reduced ICT. Furthermore, in the phenolic cyano-substituted chromophore, loss of phenolate donation and strong electron withdrawal means the HOMO is S-centred, such that overlap with LUMO, centred on the I ring/N-acylimine, is weak. Strong absorbance therefore stems from excitation from HOMO-1 (-6.50 eV) and HOMO-2 (-6.78 eV) (see Supplementary Figure S8 for frontier molecular orbital map). Under these conditions, the *trans* isomer is favoured because it maximizes residual π-overlap and stabilizes the electronic structure, whereas the *cis* form cannot compensate for geometric distortion between the P and I rings through sufficient π-conjugation. This is borne out experimentally where the P ring shifts out of plane with the I ring less in the *trans* form crystal structure compared *cis* form [11].

Incorporating a chloro-group at the 3-tyrosine position will still introduce steric bulk compared to the native mKate chromophore as chlorine will project further from the phenol ring (∼3.5 Å) than a hydrogen (∼2.3 Å) but less than the cyano group (∼4.1 Å) so acting a good intermediate group; Cl will though have a larger effective radius (∼1.75Å) than both hydrogen (∼1.2 Å) and cyano (∼1.5 Å). Incorporation of 3ClY in place of Y64 (mKate-CRO-3ClY) results in a complex absorbance spectrum (Figure 5a). Two spectral peaks are observed; a major form with a λ_max_ at 586 nm (ε=24,000 M^-1^cm^-1^) and minor form at 493 nm (ε=10,300 M^-1^ cm^-1^). The major peak is likely to represent the *cis*-phenolate form, based on TD-DFT predictions (Supplementary Table S1). λ_max_ lies either side of the DFT predicted value for the **U** (578 nm) and the **D** (598 nm) so it is difficult to ascertain if one or the other is favoured. The slight hypsochromic shift compared mKate is indicative of the electron withdrawing nature of chlorine, albeit weaker than the cyano group, which has a relatively weak inductive and resonance action. As with the *cis* phenolate form of mKate, a similar picture emerges from 3-chloro-substituted *cis* phenolate (Figure 5c), which also lowers the energy of HOMO and LUMO by the same amount, but is predicted to have slight red-shift compared to mKate. The chlorine can donate electron lone pairs to the conjugated system so the P ring retains electron density and acts an effective donor to the I ring/N-acylimine acceptor system in ICT, as is the case for mKate (Figure 3c). Bond orders for mKate as well as CN and Cl variants are reported in Supplementary Table S1, and show that both electron withdrawing groups reduce conjugation across the β-methylene bridge, and so presumably promote to *cis-trans* isomerisation. We therefore ascribe that the CN apparently exists solely in *trans* due to a combination of this reduced barrier to isomerisation and local interactions of the cyano group, assuming the *cis* form is the most likely initial state formed on chromophore maturation. S143 also does not interact with the 3-Cl group as it potentially does in the presence of 3-cyano, so will not stabilise the *trans* state and promote the phenolate *cis* state through H-bonding with the phenol hydroxyl group [11]. The minor peak doesn’t correspond to any obvious predicted peak from TD-DFT analysis. However, the excitation at this wavelength does results in fluorescence emission, with two emission peaks observed at 509 nm and 619 nm (Figure 5a). Dual peak emission was also observed for mKate-CRO-3CNY, which was ascribed to *trans-* Ph-OH from (Figure 3a). The *trans*-Ph-OH of mKate-CRO-3ClY is the closest predicted species from DFT analysis but the two differ by 32-33 nm. Excitation of the major peak results in an emission peak at 619 nm, the same as that of the secondary peak observed on excitation at the lower wavelength (Figure 5a).

## 3. Conclusion

Switchable fluorescent proteins have classically provided a means to control fluorescence emission post-maturation for super resolution imaging through chromophore *cis*-*trans* isomerisation. The space required to accommodate isomerisation can be exploited to incorporate new chemical functionalities within the chromophore [24] to tune spectral properties. Ideally, the FP chromophore should retain the phenol group as replacement of the hydroxyl group significantly blue shifts the chromophore’s absorbance [28,29]. Here we report for the first time, to our knowledge, the genetic incorporation of 3-cyano-L-tyrosine into the chromophore of the switchable RFP mKate. Incorporation of 3CNY complements existing 3-tyrosine derivative such as halides and nitrosyl, [24] and adds a potentially useful imaging tag for SRS. [37] It also provides a mean to investigate the mechanistic basis of fluorescence and switching.

We show that the 3-cyano group potentially traps the chromophore in its *trans* state through a combination of local interactions with the cyano group and lowering the barrier to *cis-trans* isomerisation. Sampling the *trans* state as its default form is very rare for FPs, even those with 3-substituted phenol rings. [24,33] While we will need to confirm sampling of the *trans* state structurally, we speculate this is partly due to the relatively strong electron withdrawing nature of the cyano group, the potentially new interactions formed by the cyano group and the prior observation that the *trans* state has a more co-planer P and I ring than the *cis* state. [11] The stabilisation of the *trans* state in mKate-CRO-3CNY could also be the reason for it retaining fluorescence emission, which is absent in the original phenolic *trans* mKate.

## 4. Methods and Materials

### 4.1. 3CNY synthesis

#### 3*-Iodo-L-tyrosinato-bicyclononylboron* (*Scheme 1*)

**Scheme 1.**
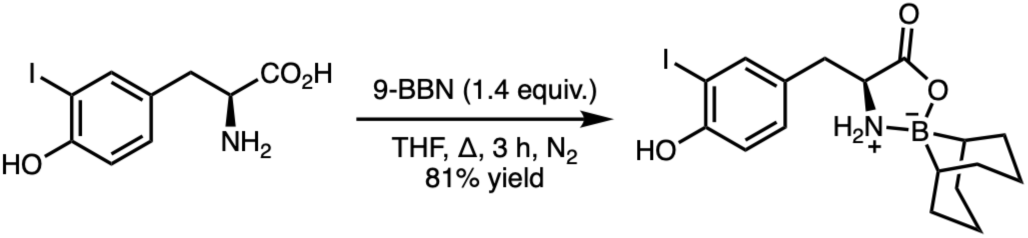

In a flame-dried round bottomed flask fitted with a condenser was added 3-Iodo-L-tyrosine (3.00 g, 9.8 mmol, 1 equiv.) and dry tetrahydrofuran (THF) (45 mL) under N_2_. The suspension was vigorously stirred for 5 min followed by the addition of 9-Borabicyclo[3.3.1]nonane (9-BBN) (0.5 M in THF, 27.4 mL, 13.7 mmol, 1.4 equiv.) slowly *via* syringe. The resulting solution was stirred under N_2_ at reflux for 3 h. The mixture was cooled and concentrated *in vacuo* yielding a yellow foam. The foam was scrapped, washed with hexanes (45 mL), filtrated and dried under vacuum to give the intended compound as an off-white solid (3.4 g, 81%), which was used directly in the next step.

#### *3-cyano-L-tyrosinato-bicyclononylboron* (Scheme 2)

**Scheme 2.**
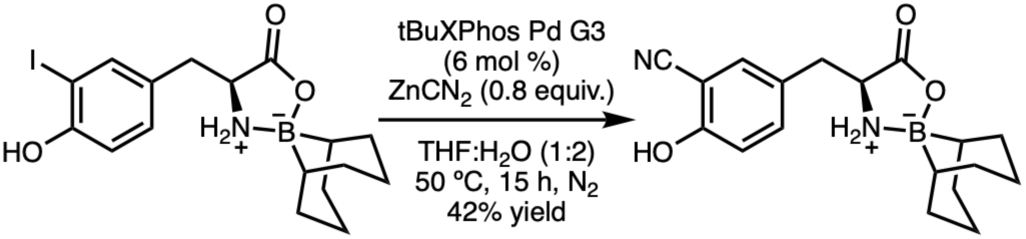

To 5 separate oven-dried 20 mL microwave vials fitted with rubber septa was added 3-iodo-L-tyrosinato-bicyclononylboron (600 mg, 1.40 mmol), 2-Di-tert-butylphosphino-2′,4′,6′-triisopropylbiphenyl (*t*BuXPhos) Pd(II) G3 (70.2 mg, 0.08 mmol, 6 mol %) and ZnCN_2_ (131 mg, 1.1 mmol, 0.8 equiv.) under N_2_. The solids were stirred under high vacuum for 15 min. Degassed THF (4 mL) and degassed water (8 mL) were added, and the resulting yellow solution was stirred for at 50 °C for 15 h. Reaction mixtures of the 5 vials were combined, cooled to rt and quenched with a saturated solution of NaHCO_3_ and diluted with ethyl acetate (EtOAc) (200 mL). The organic layer was separated, and the aqueous layer was extracted with EtOAc (2 × 200 mL). The organics were dried over MgSO_4_, filtered and concentrated *in vacuo*. Purification by column chromatography (50-70% EtOAc in hexanes, silica gel) gave 3-cyano-L-tyrosinato-bicyclononylboron as a beige solid (962 mg, 42%).

#### *3-cyano-L-tyrosine:* 3CNY (Scheme 3)

**Scheme 3.**
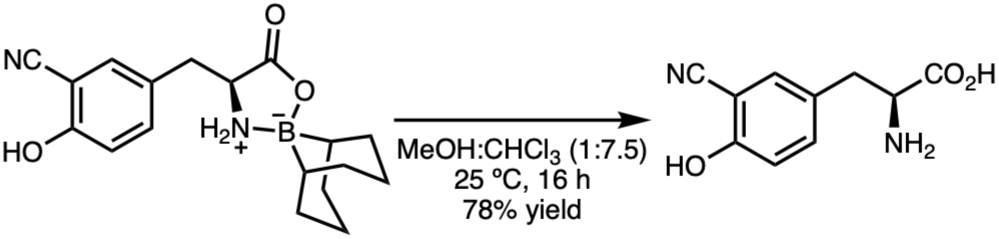

To 9 separate 20 mL microwave vials was added 3-cyano-L-tyrosinato-bicyclononylboron (106 mg, 0.32 mmol) and methanol (MeOH) (1 mL). The suspension was stirred for 30 sec and then diluted with CHCl_3_ (7.5 mL). The clear solution was stirred at room temperature for 16 h under air. The precipitated products of the 9 vials were combined, filtered, washed with CHCl_3_ and MeOH and dried under vacuum to yield the pure product as a white solid which darkens on stand (466 mg, 78% yield). **ν_max_ / cm^-1^** (thin film) 3348, 2492, 1637, 1438; **^1^H NMR (500 MHz, DMSO-*d*_6_) δ_H_** (Supplementary Figure S1) : 7.38 (d, *J* = 2.2 Hz, 1H), 7.28 (dd, *J* = 8.7, 2.3 Hz, 1H), 6.96 (d, *J* = 8.6 Hz, 1H), 3.48 (dd, *J* = 6.6, 5.3 Hz, 1H), 3.36 (d, *J* = 24.0 Hz, 1H), 2.93 (qd, *J* = 14.2, 6.0 Hz, 2H); **^13^C NMR (126 MHz, DMSO-*d*_6_) δ_C_** (Supplementary Figure S2): 169.6, 159.9, 136.0, 133.5, 127.3, 117.6, 116.8, 98.4, 55.1, 35.3; **HRMS (ES^-^)** [C_10_H_9_N_2_O_3_] requires [M^+^-H^+^] 205.0613, found 205.0613 (0.0 ppm) (Supplementary Figure S3).

### 4.2. TD-DFT and DFT analysis

Chromophore structures were taken from the available crystal structures of the low pH *trans* state (PDB 3bx9) and high pH *cis* state (PDB 3bxb). [11] The 3-cyano and 3-chloro groups were added using the Builder function in PyMOL. [59] All DFT analysis employed the Orca 6.1.0 package [60]. Geometry optimisation used the PBE-D3BJ [61,62]/def2-SVP [63] level without any constraint, and confirmed as minima using harmonic frequency calculation. Absorption spectra and frontier molecular orbitals were then calculated at optimal geometry at PBE0 [64]/def2-TZVP level in the CPCM [65] model of aqueous solution. Orbital plots were obtained using Avogadro v 1.2.0.

### 4.3. Incorporation of ncAA into mKate, recombinant protein production and purification

The gene encoding mKate was present in the pBAD plasmid and was supplied by Addgene Inc (plasmid #54826).[21] The amber stop codon was introduced in place of the Y64 codon using Q5 mutagenesis kit by New England Biolab using the forward primer 5’-CATG**TAG**GGCAGCAA-3’ and reverse primer 5’-AAGCTGGTAGCCAGG-3’ under the manufactures guidelines. 3CNY was incorporated using the 3-nitro-L-tyrosine aminoacyl-tRNA synthase-tRNA cognate pair (pDuleA7; AddGene #174078). [66] 3ClY was incorporated using the 3-chloro-L-tyrosine aminoacyl tRNA synthase-tRNA pair pEVOL plasmid from Professor Jiangyun Wang at Chinese Academy of Sciences. [23] *E. coli* TOP10 cells (Invitrogen/Thermo Fisher Scientific) were transformed with the mKate pBAD plasmid (ampicillin; working concentration 100 μg/ml) alone for native mKate production or co-transformed together with either pDuleA7 (tetracycline; working concentration 100 μg/ml) or pEVOL (chloramphenicol; working concentration 50 μg/ml) onto LB agar plates containing the appropriate antibiotics. Recombinant mKate expression was performed as described previously for other RFPs. [51] Briefly, a single colony was used to inoculate a 5 mL overnight culture, which was then used to inoculate 2xTY media supplemented with 50 μg/mL ampicillin. The cultures were left to grow at 37°C until they reached an OD600 of 0.6, when 0.2% (v/v) arabinose was added to induce expression. The cultures were left to incubate overnight at 37° in a shaking incubator. For ncAA incorporation, an autoinduction medium was used as described previously [66–68]; the media was supplemented with 1 mM of 3ClY or 0.1 mM of 3CNY. 3ClY was supplied by Fluorochem.

The production cultures were centrifuged and resuspended in 50 mM Tris HCl pH 8 and the cells lysed using a French pressure cell. Purification was performed using an AKTA Purifer using a 5 ml His Trap™ HP column (Cytiva) equilibrated in 50 mM TrisHCl pH8, 10 mM imidazole for Nickel-affinity chromatography. Bound protein was eluted by washing the column with 500 mM imidazole (pH 8.0). Pooled protein samples were then subjected to size exclusion chromatography (SEC), using a Superdex 75/300 increase 10/300 GL (Cytiva, Amersham, UK) that was equilibrated with 50 mM Tris-HCl buffer pH8.0. The purity of the proteins was then checked by SDS-PAGE. Incorporation of 3CNY was confirmed by liquid chromatography–mass spectrometry (LCMS) using a Waters Acquity UPLC/Synapt G2-Si QTOF mass spectrometer by the Mass Spectrometry facility in the School of Chemistry, Cardiff University.

### 4.4. Absorbance and fluorescence spectroscopy

Absorbance spectra were recorded using an Agilent Cary 600 spectrophotometer at 1-5 μM protein in a 1 cm path–length quartz cuvette. Protein concentrations were calculated using the known molar absorbance coefficient for mKate (45,000 M^-1^cm^-1^), which was independently verified using a Bradford protein assay. The equivalent value at 280 nm was then determined to be 46,700 M^-1^cm^-1^ and applied to the ncAA variants to determine their protein concentration via their A_280_, as has been used previously for other RFPs. [51] Emission and excitation spectra were recorded using a Varian Cary Eclipse Fluorimeter at 1-2 μM protein concentration, with the sample loaded into a 400µl quartz cuvette. Fluorescence emission spectra were recorded using a 5 × 5 mm quartz cuvette, and data were collected with a 5 nm slit width at a rate of 600 nm/min. Each protein was excited at wavelengths stated in the main text. When measured, quantum yields were determined using a comparative method described previously [68]. mCoral [51] was used as the standard for mKate-CRO-3CNY and mCherry [68] for mKate. Buffers used to record spectra at each pH were: pH 8, 50 mM TrisHCl; pH 4, 100 mM acetate buffer; pH 2, 100 mM glycine buffer.

## Supplementary Materials

The following supporting information can be downloaded at XXXX.

## Author Contributions

C.J.S. generated mutants, produced protein, undertook spectral analysis. J.J.K.M provided proof for 3CNY incorporation, generated mutants, produced protein, undertook spectral analysis. J.H., S.A.E and D.S. synthesised 3CNY. W.L and P.B. contributed to the conception of the project. J.A.P undertook DFT analysis. L.C.M contributed to 3CNY synthesis and the conception of the project and directed the project. D.D.J. contributed to project conception and directed the project, contributed to general data analysis. All authors contributed to the writing of the paper and analysing data. All authors have read and agreed to the published version of the manuscript.

## Institutional Review Board Statement

Not applicable.

## Informed Consent Statement

Not applicable.

## Data Availability Statement

All spectroscopic data will be made available via FigShare (to be released on publication). https://figshare.com/s/750d473dfc3b7744b945.

## Acknowledgments

The authors would like to thank the Cardiff School of Biosciences Protein Technology Hub for helping with the production and analysis of proteins and the Cardiff School of Chemistry Analytical services for mass and NMR spectrometry. We would like to thank Ryan Mehl, Oregon State University, for supplying pDuleA7 (via a purchase from AddGene) and Professor Jiangyun Wang, Chinese Academy of Sciences, for supplying the 3-chloro-L-tyrosine aminoacyl tRNA synthase-tRNA pair pEVOL plasmid.

