## Supplementary material for "Genetic encoding of 3-cyano-tyrosine and its use in controlling the chromophore isomeric state of the fluorescent protein mKate": SI Figures and Tables

\# These authors contributed equally.

* Correspondence:

**Supplementary Table S1. DFT analysis of mKate chromophore forms**

| mKate chromophore form | λ $S_0$-$S_1$ (nm) | HOMO (eV) | LUMO (eV) | Gap (eV) | Bond orders[a] |
|---|---|---|---|---|---|
| **Natural chromophore** | | | | | |
| *trans* Ph-OH | 455 | -6.31 | -3.35 | 2.96 | 1.53 1.06 |
| *cis* Ph-OH | 462 | -6.34 | -3.29 | 3.05 | 1.54 1.03 |
| *trans* Ph-$O^-$ | 530, 505 | -5.38 | -3.00 | 2.38 | 1.45 1.14 |
| *cis* Ph-$O^-$ | 565, 526 | -5.31 | -2.98 | 2.33 | 1.45 1.11 |
| **3CNY incorporation** | | | | | |
| *trans* 3-CN **U** Ph-OH | 446 | -6.45 | -3.45 | 3.00 | 1.55 1.04 |
| *cis* 3-CN **U** Ph-OH | 436 | -6.45 | -3.41 | 3.04 | 1.57 1.01 |
| *trans* 3-CN **U** Ph-$O^-$ | 538, 511 | -5.68 | -3.15 | 2.53 | 1.49 1.10 |
| *cis* 3-CN **U** Ph-$O^-$ | 565, 533 | -5.64 | -3.13 | 2.51 | 1.49 1.07 |
| *trans* 3-CN **D** Ph-OH | 444 | -6.44 | -3.45 | 2.99 | 1.55 1.03 |
| *cis* 3-CN **D** Ph-OH | 438 | -6.45 | -3.39 | 3.06 | 1.57 1.01 |
| *trans* 3-CN **D** Ph-$O^-$ | 537, 506 | -5.69 | -3.14 | 2.55 | 1.49 1.10 |
| *cis* 3-CN **D** Ph-$O^-$ | 564, 532 | -5.62 | -3.12 | 2.50 | 1.49 1.08 |
| **3ClY incorporation** | | | | | |
| *trans* 3-Cl **U** Ph-OH | 460 | -6.42 | -3.45 | 2.97 | 1.55 1.04 |
| *cis* 3-Cl **U** Ph-OH | 452 | -6.43 | -3.44 | 2.99 | 1.57 1.00 |
| *trans* 3-Cl **U** Ph-$O^-$ | 538 | -5.50 | -3.12 | 2.38 | 1.47 1.12 |
| *cis* 3-Cl-**U** Ph-$O^-$ | 578 | -5.43 | -3.10 | 2.33 | 1.48 1.08 |
| *trans* 3-Cl **D** Ph-OH | 461 | -6.42 | -3.45 | 2.97 | 1.55 1.04 |
| *cis* 3-Cl **D** Ph-OH | 451 | -6.43 | -3.44 | 2.99 | 1.57 1.00 |
| *trans* 3-Cl **D** Ph-$O^-$ | 538 | -5.50 | -3.12 | 2.38 | 1.47 1.11 |
| *cis* 3-Cl **D** Ph-$O^-$ | 598 | -5.29 | -3.06 | 2.23 | 1.47 1.09 |

a, Mayer bond orders for β-methylene bridge, reported in the order C(I ring)—C(methylene), C(P ring)—C(methylene)

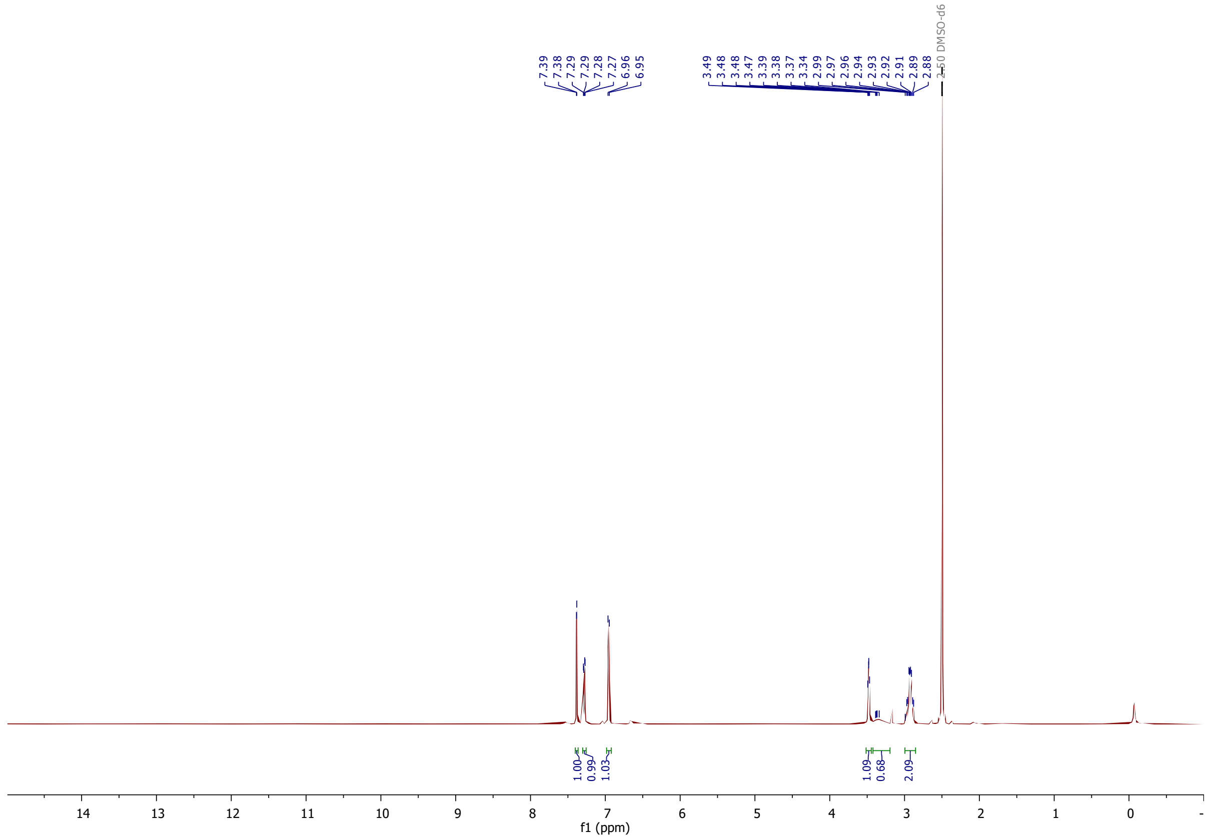

**Supplementary Figure S1.** $^{1}$H NMR (500 MHz, DMSO-d6) of 3CNY.

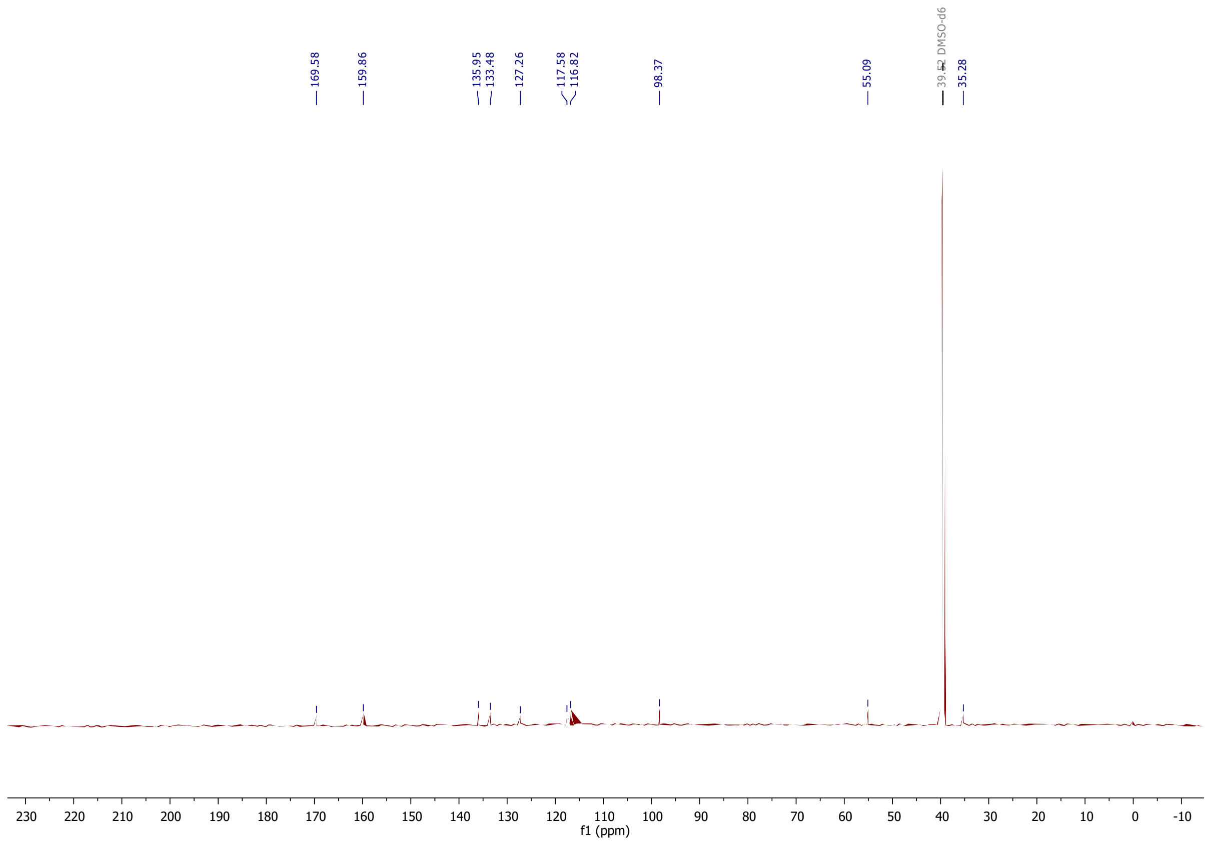

**Supplementary Figure S2.** $^{13}$C NMR (126 MHz, DMSO-$d_6$) of 3CNY

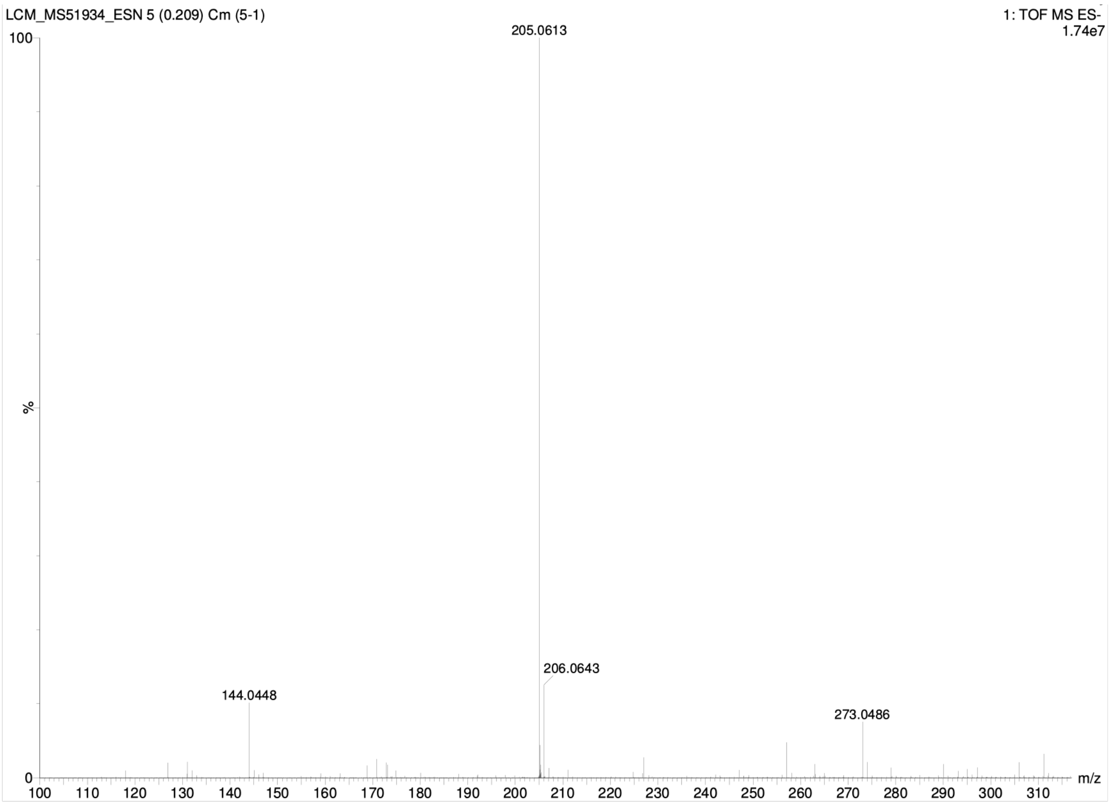

**Supplementary Figure S3.** HRMS (ES$^-$) [$C_{10}H_9N_2O_3$]. Calculated [$M^+$-$H^+$] 205.0613, measured 205.0613 (0.0 ppm).

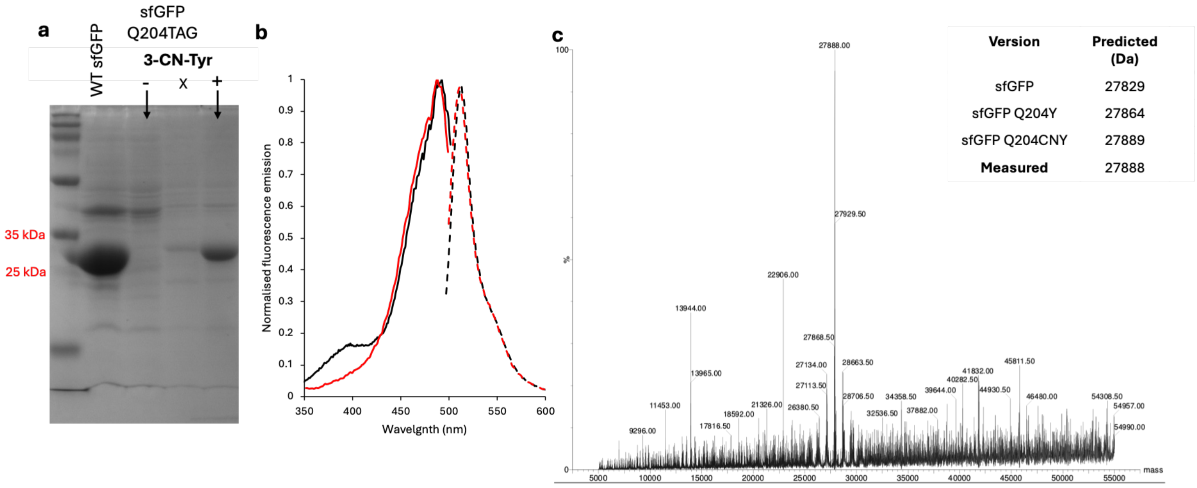

**Supplementary Figure S4. Incorporation of 3CNY in sfGFP-Q204TAG.** (a) 3CNY dependent production of sfGFP Q204CN-Y. The lane marked with an X is another ncAA (3-$NO_2$Y) analysed at the same time. (b) The whole *E. coli* cell fluorescence excitation spectra (solid lines, monitored by emission at 511 nm) and emission spectra (dashed lines, monitored on excitation at 488 nm). The WT sfGFP is coloured black and sfGFP Q204CNY coloured red. The spectra were normalised to the maximal value observed in the excitation spectra. (c) Mass spectrometry analysis of sfGFP Q204CNY. The predicted values for different versions of sfGFP Q204X are shown.

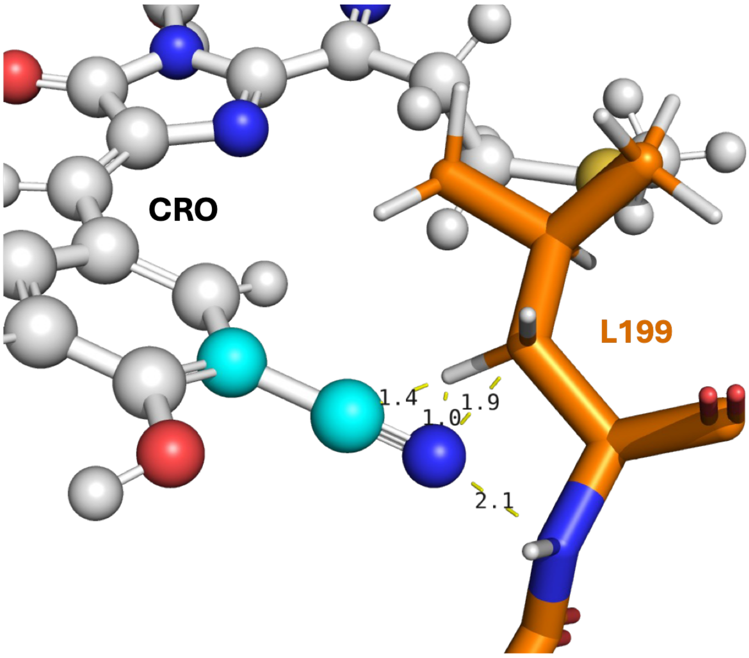

**Supplementary Figure 5.** Model of the *cis*-CNY-**D** form of the mKate chromophore. Distances (yellow lines, with distances given in Å) between the cyano group (coloured as cyano spheres) and selected L199 atoms are shown.

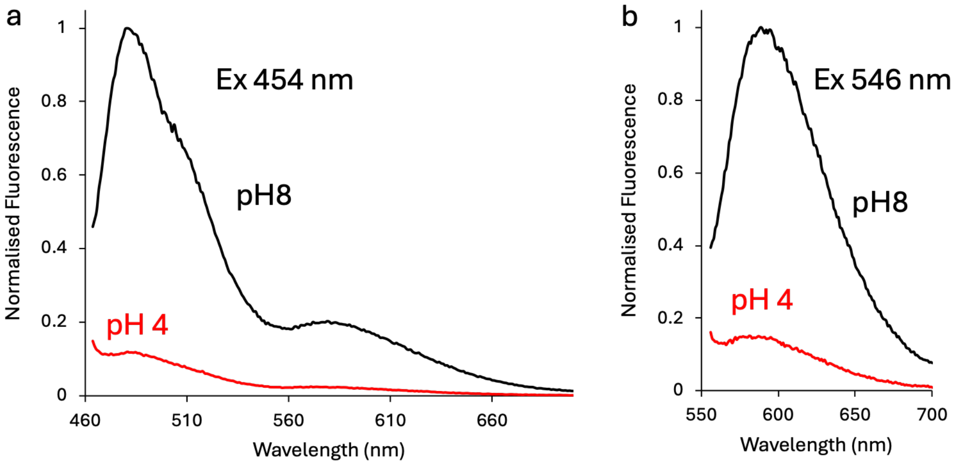

**Supplementary Figure S6.** Influence of pH on fluorescence emission spectra of mKate-CRO-CNY. Emission spectra on excitation at (a) 454 nm and (b) 546 nm). Black and red lines represent emission spectra at pH 4 and 8, respectively. Fluorescence emission is normalised to the maximum peak value at pH 8 for each excitation wavelength. Absorbance spectrum was too weak to record at pH 2.

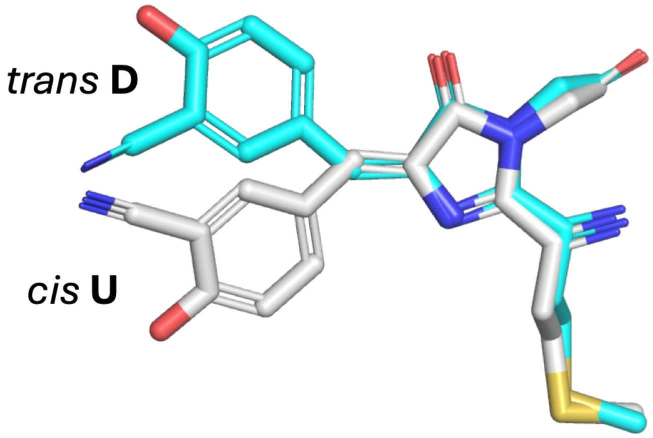

**Supplementary Figure S7.** Model of *trans* **D** (cyan) and *cis* **U** rotamer isomer forms

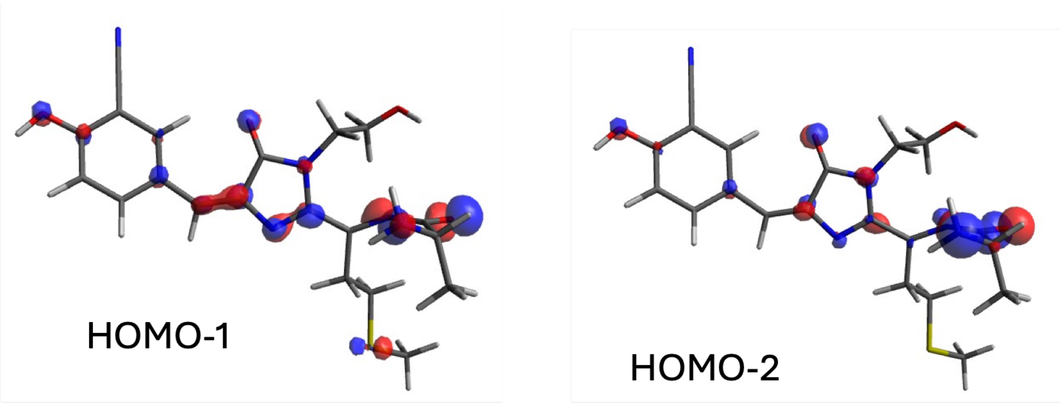

**Supplementary Figure S8.** Frontier molecular orbitals for HOMO-1 and HOMO-2 states for the phenolic *trans* mKate-CRO-3CNY.
